# A diagnostic plasma omics-biomarker for Alzheimer’s disease informed by microglial single-cell transcriptomics: A pilot study

**DOI:** 10.64898/2026.04.30.721959

**Authors:** Michael W. Lutz, Zhaohui Man, Yifei Zheng, Srilakshmi Venkatesan, Ornit Chiba-Falek

## Abstract

**Background:** The current biomarker framework for the diagnosis and staging of Alzheimer’s disease (AD) relies mainly on neuropathological features; thus, its performance for diagnosis is limited prior to the initiation of neurodegeneration. Here, we leveraged transcriptomic data to develop a new framework for omic-informed blood-based diagnostic biomarkers for AD from early-stage.

**Methods:** Microglial gene expression from single-nucleus (sn)RNA-seq data was analyzed via 6 statistical methods to identify candidate panels of genes predictive of AD. A total of 78 gene panels, 30–2000 genes in size, were selected and evaluated for their ability to distinguish AD patients from controls. Three top-ranked panels of 300, 50 and 30 genes were transferred to blood (monocyte) transcriptomic data obtained from living subjects via a graph-based mapping approach based on optimal transport statistics.

**Results:** The 300-panel method resulted in an AUC of 0.7 and moderate accuracy (75%) in classifying AD; however, the accuracy in predicting cognitively normal patients was lower (53%). While the 300 genes provided high accuracy, inspection of the distribution of p values for the gene set revealed that the panel could be greatly reduced in size to capture the most significant differences between AD patients and cognitively normal individuals. The accuracy and specificity of the 50 and 30 panels demonstrated similar AUC values but improved the balance between the prediction of AD patients and normal controls. Specifically, the 50-gene panel resulted in an AUC of 0.7, with 65% AD accuracy and 71% normal accuracy.

**Conclusions:** Integrating multiomics datasets into the AD biomarker discovery pipeline offers a powerful modality to increase precision and comprehensiveness in AD research and clinical applications.

## Introduction

The current biomarker framework for the diagnosis and staging of Alzheimer’s disease (AD), including the recent and first FDA-approved AD blood test[1, 2], relies mainly on amyloid beta (Aβ) and tau (primarily p-tau fragments) pathologies as core markers. Thus, their performance is limited, particularly prior to the beginning of neurodegeneration and atrophy. While assays based on positron emission tomography (PET) imaging and cerebrospinal fluid (CSF) are invasive, expensive and less accessible, newer plasma tests reduce these limitations and increase diagnostic accessibility[1, 2].

AD initiates decades before clinical symptoms and neuropathological hallmarks appear. The new AD blood test, which is based on the pTau217/ß-amyloid 1-42 plasma ratio, can predict the presence of amyloid plaques and is intended for patients who are cognitively impaired. Thus, the accuracy and sensitivity of the available plasma tests are limited, especially in presymptomatic stages, highlighting the unmet clinical need for new diagnostic biomarkers enabling precise early diagnosis. Furthermore, predictive biomarkers to identify subjects at high risk of developing AD years and even decades before we notice symptoms would be beneficial for the design of the next generation of AD clinical trials; therefore, these biomarkers are crucial for shifting toward prevention treatments or disease-modifying therapies (DMTs) prior to the initiation of severe neurodegeneration processes, including the accumulation of beta amyloid and tau in the brain.

The role of neuroinflammation and the involvement of microglia in the pathogenesis of neurodegenerative diseases, including AD and related dementias, have been well established[3–8]. Genome-wide association studies (GWASs) have shown that microglia-expressed genes are involved in AD risk, and transcriptomic studies have demonstrated dysregulation of gene expression in microglia from the postmortem brains of patients compared with those from age-and sex-matched controls[9–13]. Moreover, it has been suggested that microglia constitute one of the earliest cell subtypes showing gene expression changes in AD[14–17]. We aim to translate this knowledge into the development of accurate, sensitive and noninvasive diagnostic biomarkers for presymptomatic AD.

In this study, we developed a new biomarker framework and represented the first stage in the development pipeline of a transcriptomic-based AD blood-biomarker for living individuals (**Fig. 1**). First, we identified transcriptomic biomarkers based on AD-associated microglial differentially expressed genes (DEGs) using a publicly available single-nucleus (sn)RNA-seq dataset [18, 19]. We then applied a label transfer approach to translate the discovered microglial DEG-signatures into monocyte samples, both cell-types of the myeloid lineage. Finally, we evaluated performance of the microglial transcriptomic signatures to assign phenotypic categories in the monocyte samples. Our approach is novel in the field blood biomarker discovery for AD diagnosis even in early stages, as the biomarker derived from brain microglia rather than by direct examination of the blood cells *e.g.,* monocytes. This is a promising strategy because it captures a well-characterized core brain molecular signature associated with immune and inflammatory processes in microglia, an established disease-driving cell type that mediates pathogenic mechanisms contributing to AD onset.

**Figure 1.**
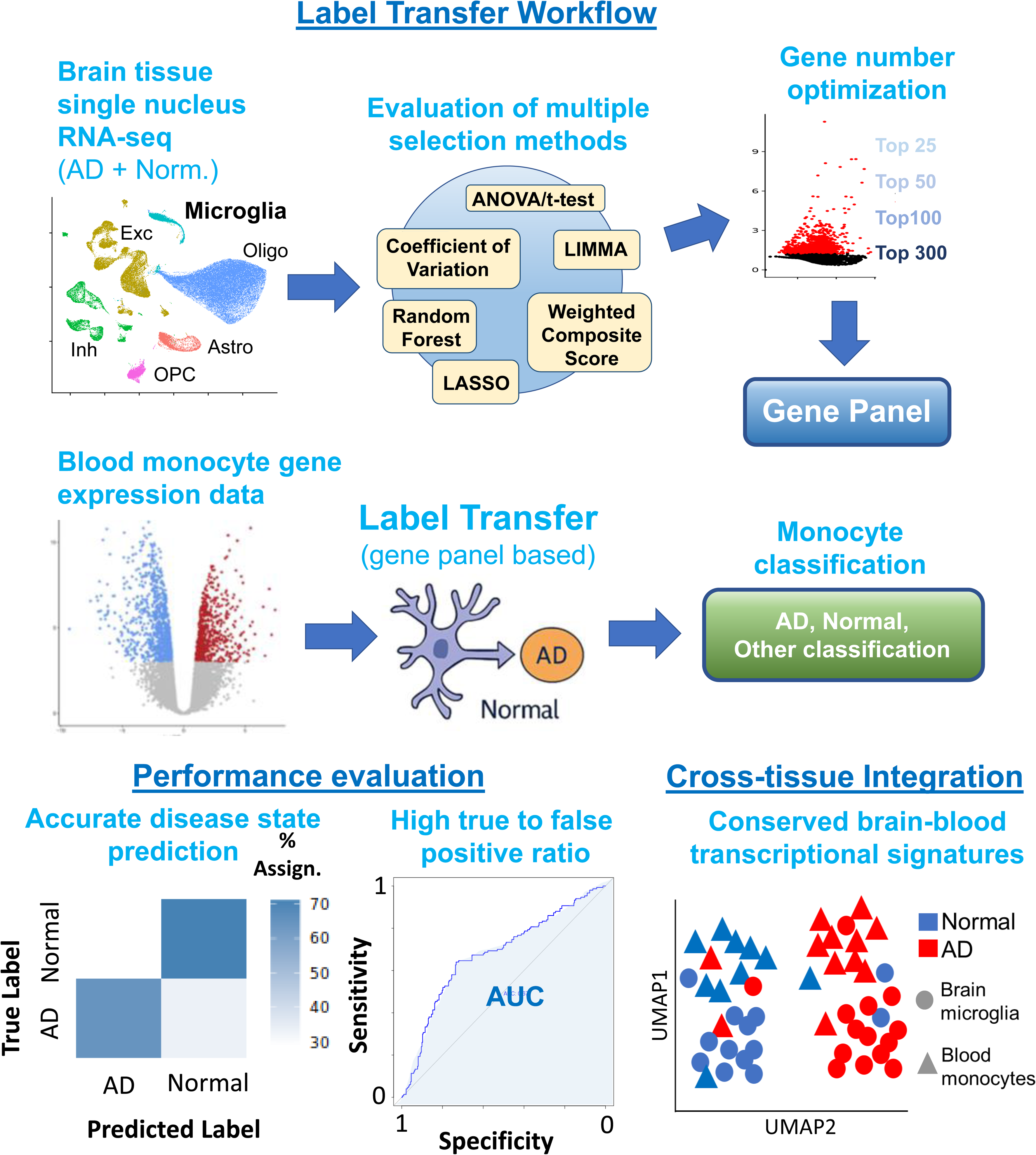
The workflow for the development of plasma transcriptomic biomarkers for AD prediction on the basis of the microglial transcriptome.

## Methods

### Study design and data sources

We developed a bioinformatics strategy to evaluate the transcriptional signatures (gene panels) of Alzheimer’s disease in brain microglia that could be transferred to peripheral blood monocytes for diagnostic applications. snRNA-seq data from microglia were obtained from the Religious Orders Study and Memory and Aging Project (ROSMAP) cohort via the Alzheimer’s Disease Knowledge Portal (syn31512863). These samples were derived from the dorsolateral prefrontal cortex tissue of participants with well-characterized cognitive status. Notably, samples from individuals with AD were, on average, 2.4 years older than those from individuals with normal cognition (*P*<0.0001). Peripheral blood monocyte (PBMC) bulk RNA-seq data were obtained from the same research program (syn22024498) to serve as the target domain for label transfer; all PBMC samples were obtained pre-mortem. The PBMC samples were obtained an average of 1.6 years (SD=2.7) before death. Descriptive statistics for clinical and demographic variables are provided in **Table 1**. One of the study aims was to examine predictive accuracy in monocyte samples that were independent from the microglial samples. There were 81 of the 236 monocytes samples that overlapped with the microglial samples, the remainder of the samples (n=155) were unique; all statistical analyses accounted for overlapping samples by calculating predictive accuracy for the non-overlapping samples.

**Table 1.** Clinical and demographic variables for ROSMAP microglial and blood monocyte snRNA-seq samples.

|  | Microglia |  | Monocytes |  |
| --- | --- | --- | --- | --- |
| Parameter | AD | NCI | AD | NCI |
| N | 145 | 125 | 121 | 115 |
| age at death | 88.2<br>(3.4) | 85.8<br>(5.1) | 86.8<br>(4.2) | 84.9<br>(5.5) |
| N, % female | 106,<br>73% | 80,<br>64% | 78,<br>64% | 81, 70% |
| APOE alleles |  |  |  |  |
| 0 | 87, 60% | 100,<br>80% | 77,<br>65% | 91, 81% |
| 1 | 53, 37% | 25,<br>20% | 39,<br>33% | 21, 19% |
| 2 | 4, 3% | 0, 0% | 3, 2% | 0, 0% |
| Braak score | 4.1 (1.0) | 3.1<br>(1.2) | 4.2<br>(1.0) | 3 (1.2) |
| Cerad score | 1.7 (0.9) | 2.7<br>(1.2) | 1.8<br>(1.0) | 2.8 (1.2) |
| PMI | 7.3 (4.3) | 7.3<br>(4.3) | 7.9<br>(4.4) | 9.5 (8.0) |

For validation, the prediction of MCI was tested in an independent cohort, data were obtained from the Mayo Clinic study of Aging obtained from the Alzheimer’s Disease Knowledge Portal (syn22024998), descriptive statistics are provided in **Supplemental Table S1**.

Details on the cohort composition, quality control procedures and transcriptomic data processing are provided in **Supplementary Methods**.

In contrast to the PBMC samples, these samples are cell-free bulk RNA (cfRNA) from PAXgene blood tubes. While both label transfer performance evaluations provide statistics on predictive accuracy for disease diagnosis from the microglial transcriptome, the PBMC sample results are focused specifically on immune cell activity for later-stage AD while the cfRNA samples are centered on translation from the brain microglial signatures to earlier stage disease (MCI) using samples that would be more easily obtained during a clinical trial.

### Biomarker gene panel construction

We employed a systematic approach to identify optimal gene signatures by testing six complementary statistical feature selection strategies on the microglial training data. Statistical details are described in **Supplementary Methods**.

### Deep Joint Distribution Optimal Transport architecture

Cross-tissue label transfer uses a deep joint distribution optimal transport (DeepJDOT) neural network framework designed to align feature distributions between the source and target domains while preserving class-discriminative information. The architecture consisted of a shared encoder network with two fully connected layers (input → 2×hidden → hidden dimensions), incorporating batch normalization and dropout regularization (rates of 0.2--0.4) to prevent overfitting. Details of this approach, specifically optimization process, are in **Supplemental Methods**.

### Label transfer evaluation and validation

Statistical Details on the evaluation of the label transfer approach including ROC analysis, UMAP visualization, quantification of spatial distributions of cell types and disease states and biological pathway analysis are in **Supplemental Methods**.

## Results

### Selection of gene panels of differentially expressed genes in microglia in AD

The first step identified high-resolution microglial transcriptomic signatures for distinguishing between individuals with and without AD. We obtained snRNA-seq data from the dorsolateral prefrontal cortex (DLPFC) of the ROSMAP cohort, which consisted of 145 samples with a cognitive diagnosis of AD and 125 samples with a diagnosis of normal cognition[18, 19] (**Table 1**). The analysis utilized the microglial cell pseudobulk RNA-seq data extracted from this DLPFC snRNA-seq dataset and applied six different gene selection statistical methods (**Fig. 2, Extended Data Supplement**). Noteworthy, our strategy was supported by prior publications reporting microglial differential gene expression in the analysis of ROSMAP snRNA-seq data [20, 21].

**Figure 2.**
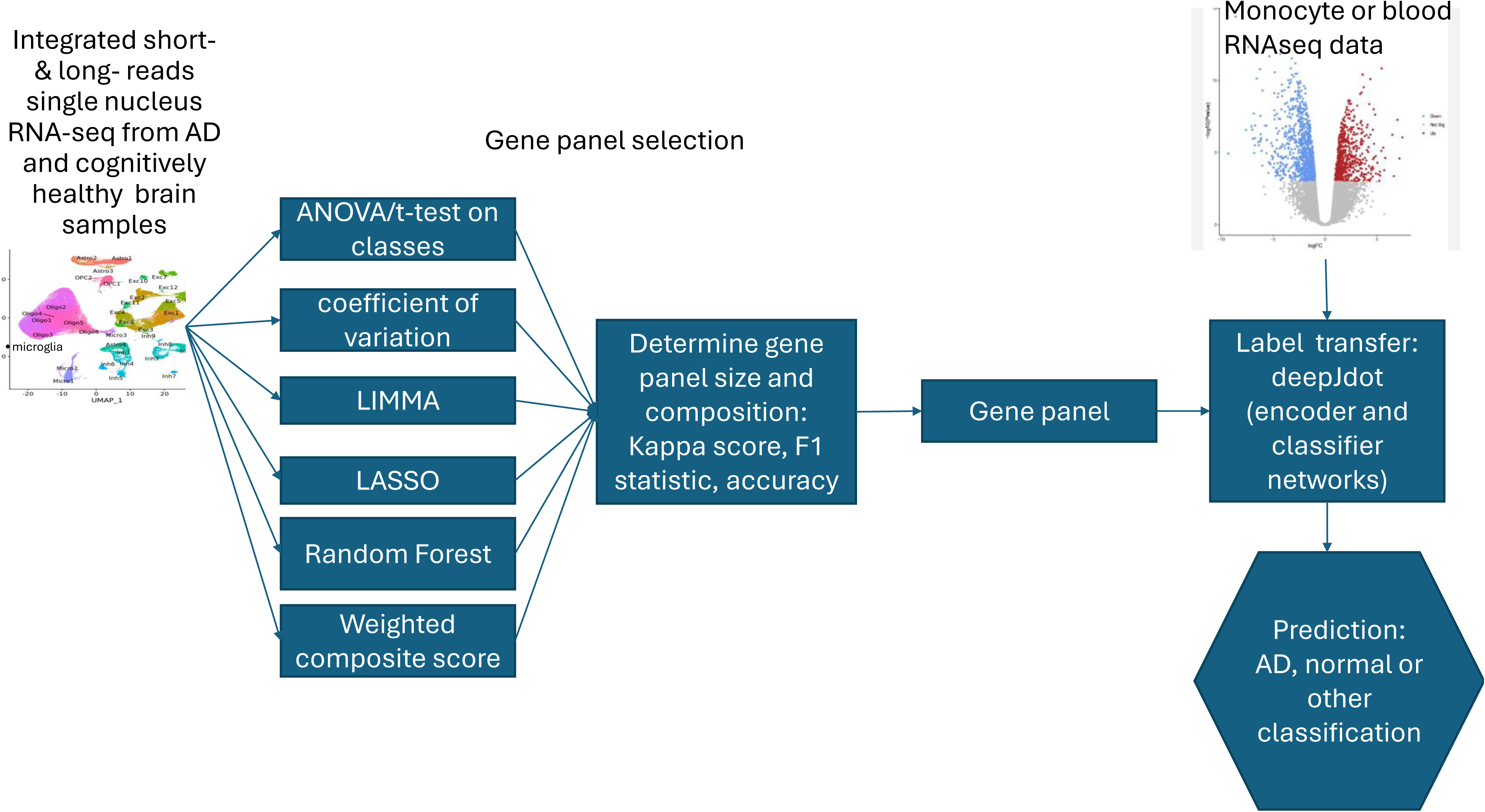
Overview of the statistical methods applied in the biomarker discovery pipeline: selection of gene panels on the basis of single-nucleus data from microglia, label transfer from microglia to monocytes, and performance evaluation in monocytes.

### Evaluation of the accuracy and precision of the different microglial gene panels in the classification of AD diagnosis

Next, we evaluated the performance measures of each gene panel (total 78) to predict AD diagnosis vs normal cognition (**Fig. 3**). The performance measures included accuracy (true positives + true negatives)/total predictions), (**Fig. 3A)**, precision (true positives/(true positives + false positives)), (**Fig. 3B)**, recall, i.e., the proportion of actual positives (AD cases) that were correctly identified or (true positives/(true positives + false negatives)), (**Fig. 3C)**, and the F1 score, the harmonic mean of precision and recall or ((2·true positives)/(2·true positives + false positives + false negatives)) (**Fig. 3D**). Overall, the 30-gene panel (**Supplemental Table S2**) selected with the combined algorithm and the 50-gene panel (**Supplemental Table S2**) based on the t test showed the best performance as demonstrated by F1 scores (**Supplemental Table S3**). In addition, we examined 6 larger panels of genes (100--2000), and a panel of 300 genes selected by the limma algorithm (**Supplemental Table S2**) demonstrated the best performance measures (**Supplemental Table S3**). Comparisons between statistical methods for selection and gene panel sizes and specific performance measures are discussed in the **Extended Data Supplement**.

**Figure 3.**
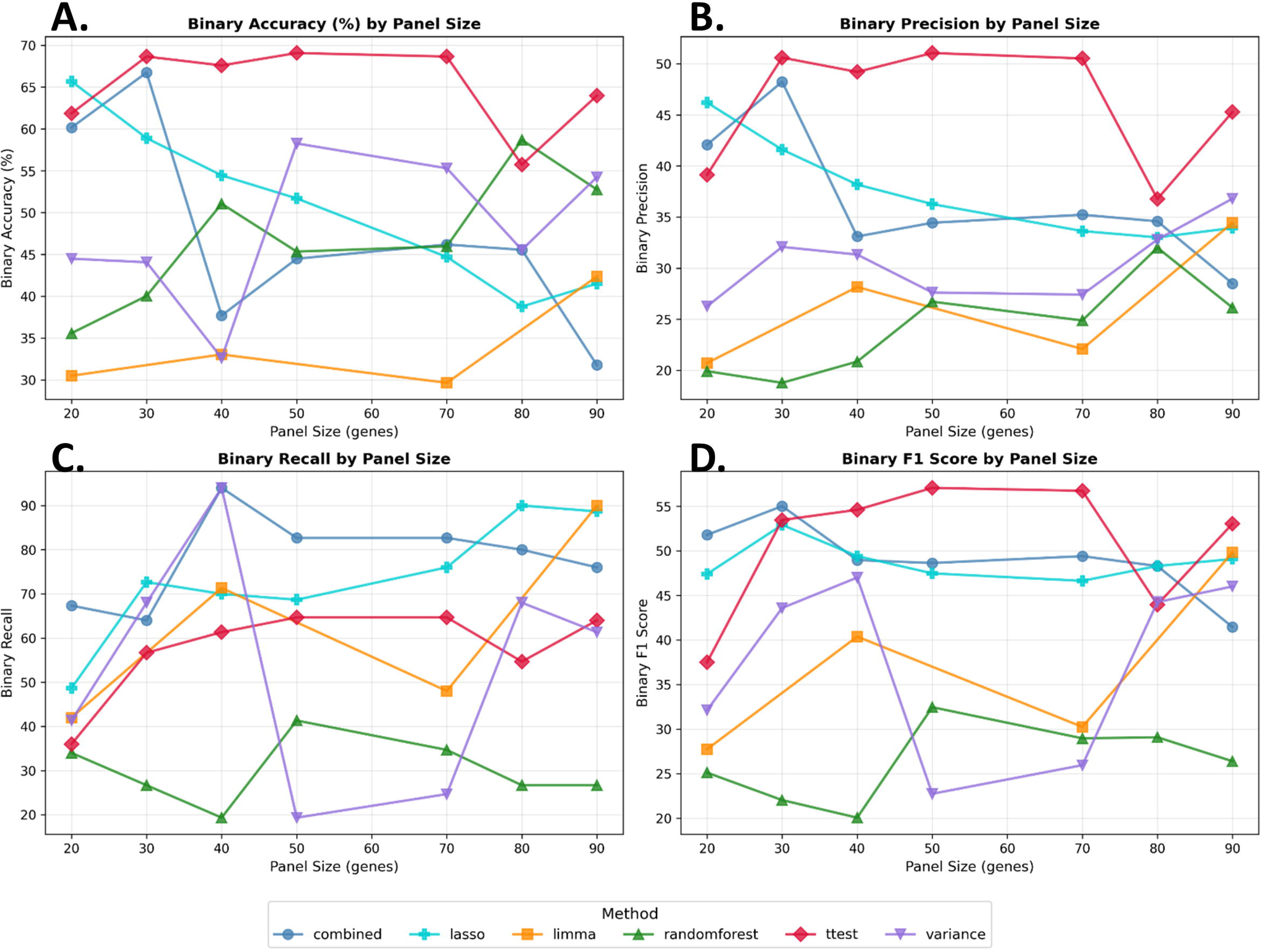
Performance measures for each of the six gene selection methods. Each plot shows a statistical measure of the performance of the gene panel differentiating AD cases from controls for panels of different sizes. (**A**) Accuracy. (**B**) Precision. (**C**) Recall. (**D**) F1 score.

### Evaluation of the selected gene panels to classify AD vs. normal subjects in monocyte transcriptomic data

In the second step, we used monocyte RNA-seq data[22, 23] (**Table 1**) to test the label transfer performance of three selected panels of 300, 50 and 30 genes (**Supplemental Table S2**) and their sensitivity, specificity and area under the curve (AUC) to classify AD vs normal cognition.

The panel of 300 genes selected by the limma algorithm had an AUC of 0.7 for label transfer to monocytes, with strong accuracy in predicting the AD class (75%). However, the accuracy in the prediction of cognitively normal individuals was lower at 53%. While the 300-gene panel provided high accuracy, inspection of the p value distribution for this gene panel (**Supplementary Figure 1**) revealed that the gene panel size could be greatly reduced such that it would capture the most significant differences between the diagnostic criteria. Indeed, when a p value threshold of 0.05 (-log_10_(0.05) = 1.3) was used, the number of genes included in this panel was reduced to only 8 genes.

Gene panels of 30 and 50 genes selected because of the optimal F1 score are potentially more feasible for implementation in clinical settings as transcriptomic biomarkers.

The accuracy and specificity of the 30-gene panel selected by the combined algorithm and the 50-gene panel selected by the t test in the label transfer analysis of the monocyte RNA-seq data demonstrated a good balance between the predictions of the AD and normal classes. As shown by the ROC curves, the 50-gene panel had an AUC of 0.7 (**Fig. 4A**), 65% AD accuracy, and 71% normal class accuracy, and the 30-gene panel had an AUC of 0.7 (**Fig. 4B**), 64% AD accuracy and 68% normal class accuracy.

**Figure 4.**
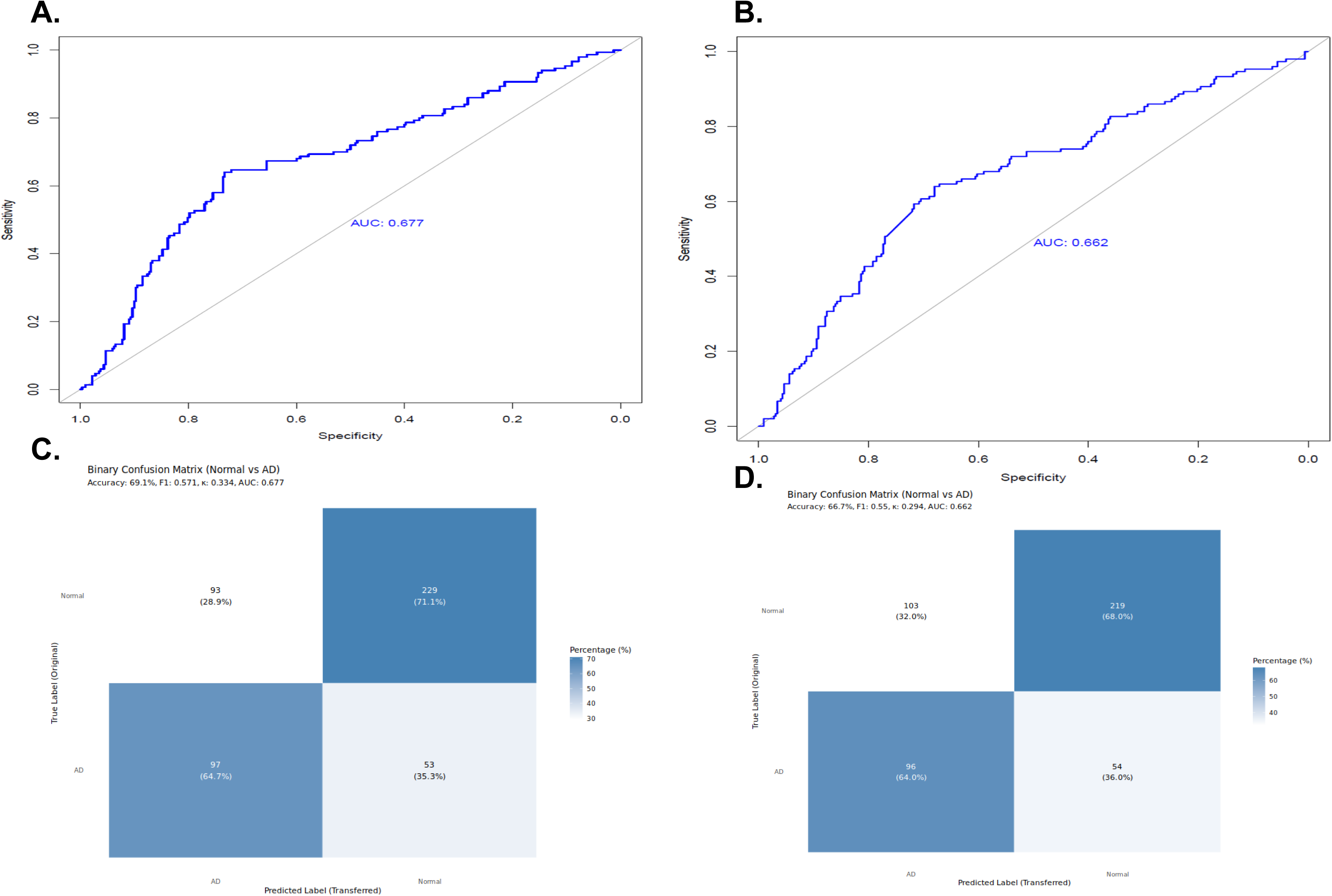
Performance of the microglial-based gene panels predicting AD from monocyte data. (**A**) ROC curve for the 50 gene panel, t-test selection algorithm, (**B**) ROC curve for the 30-gene panel, combined algorithm for selection. (**C**) Confusion matrix of accuracy values for the 50-gene panel, t-test selection algorithm. (**D**) Confusion matrix of accuracy values for the 30-gene panel, combined selection algorithm.

Integrated UMAP diagrams revealed strong clustering by disease for both the microglial samples and the monocyte samples according to UMAP1 scores (**Fig. 5**). The 50-gene panel, which was grouped into two distinct clusters of AD samples for monocytes and one cluster for microglia, was largely separated by UMAP2 values (**Fig 5A**). In the two AD monocyte clusters, there were only a few normal microglial samples; however, in the AD microglial cluster, we observed some overlap between the normal and AD samples. The normal samples presented one cluster of monocyte samples and one cluster of microglial samples, which were primarily separated by UMAP2 values. The normal monocyte cluster covered a large range of UMAP2 values, with few misclassified AD microglia or monocyte samples. The normal microglial cluster contained predominantly normal samples, with only a few AD samples. The 30-gene panel revealed that most of the microglia and monocyte samples were in one large cluster (**Fig. 5B**).

**Figure 5.**
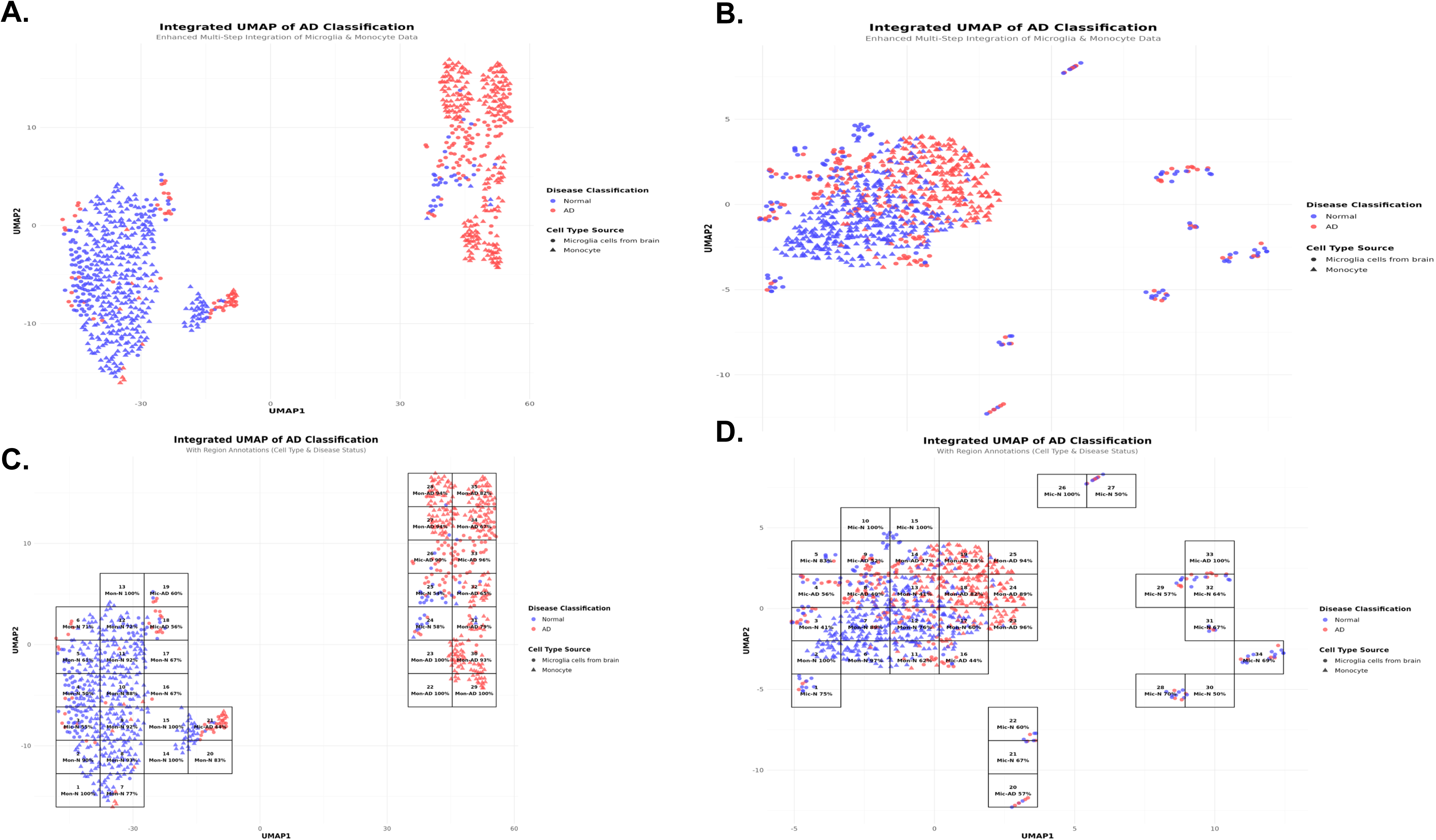
Integrated UMAP plots showing clustering of microglial and monocyte data for samples from AD and cognitively normal individuals. UMAP visualization illustrates a low-dimensional graph of the high dimensional gene expression data. (**A**) UMAP for the 50-gene panel, t-test selection algorithm. (**B**) UMAP for the 30-gene panel selected with the combined algorithm. (**C**) Cluster mapping and calculations of separation by cell-subtype and disease state for the 50-gene panel, t-test selection algorithm. (**D**) Cluster mapping and calculations of separation by cell-subtype and disease state for the 30-gene panel selected by the combined algorithm.

AD microglial and monocyte samples were separable from normal samples in terms of UMAP1 and UMAP2 values but to a lesser extent than the 50-gene panel (**Fig. 5B**).

Regional composition analysis of the UMAP clusters was performed by overlaying a 10x10 grid on the UMAP plots and counting the number of samples labeled by cell type and disease state. The 30 gene panel annotations revealed significant intermingling between cell types and disease states (**Fig. 5C**). There were distinct monocyte-enriched (left cluster) and microglia-enriched (upper/right cluster) regions, but many regions contained both cell subtypes (regions 3, 8, 9, 13, 14, and 16). Disease state (AD/normal) classification was shown primarily within monocyte populations (regions 18, 19, 23, 24, and 25 showed >85% monocyte–AD purity). However, mixed regions, including region 13 (41% monocyte-normal, 39% monocyte-AD), were indicative that the separation by disease state was limited. In contrast, the 50-gene panel regional composition analysis revealed strong distinct separation by cell type and disease state (**Fig. 5D**). For the cell type, there was clear stratification by UMAP1 values, with monocyte-enriched regions (1, 2, 6--17, 20) showing tight clustering and microglia-enriched/mixed regions (18--26, 33) on the right and dramatically reduced intermingling, in contrast to the 30 gene panel results. For disease status stratification, the monocyte-AD regions (27--32, 34, 35) formed a compact, highly pure cluster (>93% monocyte-AD purity in regions 27--30, 35), whereas the monocyte-normal regions (1--17) were similarly compact and pure, indicating superior AD/normal discrimination in the monocyte cell type. There were fewer regions with mixed compositions than with the 30-gene panel, suggesting more effective feature-based separation of both the cell type and disease status simultaneously.

The 50-gene panel demonstrated substantially improved AD vs normal classification within monocytes; the 30-gene panel results revealed that 8/17 monocyte-containing regions (47%) achieved >80% AD or normal purity, whereas the 50-gene panel revealed that 17/32 monocyte-containing regions (53%) achieved >80% purity. Notably, the 50-gene panel resulted in highly enriched monocyte AD clusters (regions 27--35) with 93--100% purity, whereas the 30-gene panel resulted in fragmented AD classification (regions 18--25 with 46--95% heterogeneity).

Collectively, the results demonstrated that disease status determined clustering across the cell types. The multiple clusters for microglial AD samples suggested that the sample cohort consisted of several AD molecular subtypes. The UMAP plots, most notably for the 50 genes panel, showed strong separation between AD and cognitively normal samples with clusters containing both microglial and monocyte cell subtypes, pointing to conserved transcriptional signatures between the brain and peripheral monocytes, thus supporting our biomarker development strategy.

We compared the gene panels with the results previously reported for snRNA-seq analysis of PBMCs [24]. These AD-associated DEG sets were significant for 1, 8 and 71 genes (FDR<0.05) from our 30, 50 and 300 gene panels, respectively. The cell subtypes most represented for these DEGs included CD4+, CD8+ and CD56 NK cells.

### Evaluation of the selected microglial transcriptomic signatures to classify MCI vs. normal subjects

We tested the performance of the selected microglial DEG-panels as biomarkers for MCI prediction via an independent dataset, the Mayo Clinic Study of Aging (MCSA). This dataset consists of blood samples from diagnoses of cognitively normal individuals and MCI patients. In this analysis we used a balanced sample of 44 MCI patients and 44 cognitively normal individuals selected at random out of 374 samples (**Supplemental Table S1**). Consistent with the results of the AD diagnosis analysis, the 30-gene panel selected via the combined (composite score) algorithm presented the best performance for panels with <100 genes, demonstrating an optimal balance between the accuracies to predict both MCI and normal cognition. The prediction of MCI patients vs cognitively normal controls resulted in an AUC of 0.52, with the accuracy of normal class prediction showing the best performance of 64% accuracy, whereas the prediction of MCI patients was modest at 41% accuracy. The 50-gene panel selected with the t test showed an improved AUC of 0.60, with 66% accuracy in detecting MCI; however, the accuracy in detecting the cognitively normal cases decreased to 18% compared with that of the 30-gene panel. The larger gene panels of 600 genes selected by maximum variance resulted in an AUC of 0.63, with 91% accuracy in detecting the cognitively normal cases and 34% accuracy in detecting MCI. Collectively, the results of this MCI prediction analysis supported the premise that the discovered transcriptomic panels enabled the detection of early-stage cognitive decline associated with AD. Remarkably, the substantially improved performance of the larger gene panel relative to the smaller panels indicated that various transcriptomic signatures may offer different predictive accuracies depending on the stage of the disease (i.e., pre-clinical, early, late) through the course of AD development.

### Biochemical pathway analysis and gene panel overlap with AD GWAS

The network of enriched biological pathways (**Supplementary Figure 2**) contained a set of highly connected pathways relevant to neuronal processes and neurodegeneration (**Supplementary Figure 3**). For the 300-gene panel, we identified 13 AD GWAS genes These genes are involved in multiple signaling pathways, including RHO GTPases, RAS GTPases, and MAP kinases. Further details are in **Extended Data Supplement**.

## Discussion

This study is the first step in a novel framework for AD biomarker discovery informed by microglial transcriptomic signatures and based on blood-test. Here, we identified microglial transcriptomic-informed biomarkers and evaluated their performance to diagnose AD at both early and progressed disease-stages by leveraging on microglial and monocyte RNA-seq datasets, respectively. The results of our pilot study demonstrated the strong *translational* potential of transcriptomic-biomarkers as the new generation of blood-based diagnostic biomarkers for AD at early stages in living subjects.

The strategy to develop blood-based biomarkers for early-stage AD diagnosis via microglial transcriptomic data is supported by the following rationale: (1) Microglia play a key role in the etiology of AD, including the contributions of neuroinflammation and immunomodulation pathways to AD pathogenesis[3, 20]. Specifically, disease-associated microglia (DAMs) have been identified in several groups and characterized by enhanced phagocytic activity and lipid metabolism changes associated with the disease state, localization around amyloid beta plaques, and increased expression of *TREM2* and *APOE* [25–27]. (2) snRNA-seq studies identified multiple DEGs in AD microglial cell types and subtypes[9, 28–30], and microglial transcriptomic signatures of AD have been reported [5, 8, 20]. For example, compared with non-AD brains, 643 deregulated genes were identified in human brains with early amyloid pathology. (3) Microglia are involved in the early stages of AD development. (4) Microglia and peripheral blood mononuclear cells (PBMCs) are both part of the myeloid lineage. Microglia are brain immune cells that are derived primarily from primitive macrophages and can be replenished from circulating monocytes (a type of PBMC) throughout life. Furthermore, these genes share gene expression profiles. Our results revealed some overlap between the genes in the biomarker panels and AD GWAS genes, with the 300-gene panel enriched for genes involved in endocytosis, RNA splicing and neuron projection development, which are all critical pathways involved in Alzheimer’s disease. Overall, the results of this study support the strategic plan for biomarker development. Specifically, UMAP analysis revealed clusters that overlapped monocyte and microglia cell populations and were stratified by disease status, confirming the concept of deducing the disease state across these cell types. Furthermore, pathway analysis revealed that biological processes are likely affected in early-stage AD, suggesting their utility for early-stage diagnosis.

The accuracy of our selected gene panels in predicting AD in monocyte samples should be considered in the context of previous blood-based AD biomarker development studies. The performance of six AD blood biomarkers for incident all-cause and AD dementia showed strong predictive performance of 77.5% to 82.6% for p-tau181, p-tau217, NfL and GFAP, with high negative predictive values >90% but low positive predictive values (12–34%)[31]. Notably, for the prediction of AD dementia, the AUCs ranged from 0.71 for NfL to 0.77 for p-tau217[31]. Studies of blood-based gene expression data have identified genes related to AD and their ability to predict early AD[32, 33]. For the external validation experiments (cross-cohort), the AUCs for the prediction of AD were on the order of 0.70 to 0.66[32], and the Alzheimer’s Disease Neuroimaging Initiative (ADNI) was used as the input to predict AddNeuroMed1 (ANM1)[32]. Using 1604 DEGs from ANM1 for training yielded AUCs of 0.62 and 0.79 for AD prediction in the ADNI and AddNeuroMed2 cohorts, respectively[32]. Integrative analysis of single-cell and cell-free plasma RNA transcriptomics was used to derive a 340-gene panel for the prediction of AD, with reported AUCs ranging from 0.58 to 0.89 in different validation sets, via several prediction algorithms, including random forests, support vector machines and logistic regression[34]. A study of plasma proteomics identified panels of proteins that differentiate AD samples from non-AD samples with AUC values of 0.93[35, 36], although these proteins were not specifically linked to markers of neuroinflammation. An evaluation of our 30- and 50-gene panels of monocytes revealed approximately 70% accuracy in detecting disease (AD and MCI) in an independent cohort, which was comparable to the findings of previous studies. Thus, these two panels are strong candidates as blood-based transcriptomic biomarkers for AD diagnosis. These results provide the *proof-of-concept* needed for advancing the development of the identified transcriptomic biomarkers to the next stage in the pipeline, including validation studies using blood samples from living subjects.

AD is a heterogeneous disease manifested by clinical comorbidities, diverse rates of progression, ages of onset, and co-pathologies[37]. The heterogeneity of AD poses challenges to the pipeline of AD biomarker discovery. Moreover, recent multiomics studies, including cerebrospinal fluid (CSF) proteomics[38, 39], brain transcriptomics[40], and multiomic integration[41], have identified molecular subtypes of AD. The characterization of multimodal molecular subtypes of AD has provided the premise for identifying AD biomarkers via multiomic data integration[42]. While the present study focused on binary classification for AD to develop a gene panel based on fewer than 50 genes, the approach could be generalized to more strata, including phenotype-based subgroups, that reflect the heterogeneity of AD.

A recent study utilized an optimal transfer approach to map differentially expressed genes (DEGs) from ROSMAP monocyte data to peripheral blood mononuclear cell (PBMC) samples from two independent cohorts, ADNI and ANMerge[22]. This work aimed to predict AD disease subtypes and molecular signatures from blood samples from living individuals based on precise neuropathological diagnoses of postmortem brains. Similar to our study, the authors used a single cell-type based on postmortem neuropathological staging or diagnosis to perform a label transfer for the diagnosis of living patients. However, there are major differences between our study and that of Huang et al.[22]: (1) We based the label transfer process on specific, microglial transcriptomic signatures from the brain to identify an optimal statistical method for developing predictive gene panels for blood-based biomarkers, whereas Huang et al. transferred subtype labels from ROSMAP monocyte samples to peripheral blood mononuclear cell (PBMC) samples within the ADNI and ANMerge cohorts; (2) we used the diagnostic labels of normal cognition, MCI and AD, whereas Huang et al. utilized labels of control, asymptomatic AD, typical AD and low-NFT AD; (3) Huang reported 40% to 80% label transfer accuracy for the blood samples: using their optimal transport approach[22, 43] resulted in the highest accuracy (80%); Ingest[44], the Monocle[45] and MMD-MA[46] approaches yielded lower accuracy values. The accuracy of our approach using deep joint distribution optimal transport was approximately 70%, which is close to the higher accuracies reported by Huang et al.[22].

The study design was rigorous; however, several limitations should be noted. First, the ability to predict AD and/or MCI status warrants replication in other independent cohorts.

Nonetheless, the independent MCSA cohort provided valuable information related to the prediction of MCI as an indication of early disease stages. The construction of the gene panel was based on microglial cell types as a whole and did not consider the granulated subtypes of microglia, such as activated microglia and/or DAMs. It is possible that microglia signatures could be perturbed by other inflammatory processes related to neurodegeneration but not specific to AD. Additionally, although sample sizes had adequate statistical power to detect moderate effect sizes, larger sample sizes would enable the construction of alternative biomarker panels and the testing of panels with smaller effect sizes.

## Conclusions

This study represents a new frontier in AD biomarker discovery by integrating single-cell omics data, such as transcriptomic data, into the pipeline. The outcomes provide a *proof-of-concept* for continuing the development of the 30 and 50 gene panels as transcriptomic biomarkers for AD administered via a blood test. Further development of our investigational new omic-based biomarker will enable accurate and sensitive diagnosis of AD at early presymptomatic stages before the clinical onset of symptoms. Moreover, it is designed to be a blood test and is thus noninvasive and accessible. Here we showed the diagnostic utility of omics-biomarkers in clinical settings. In addition, our new AD-biomarker discovery framework could be empowered for the development of additional omics-biomarkers for other intended uses, including predicting disease risk, disease trajectories/prognosis, monitoring treatment effects, etc (REF). These omics-biomarkers accurate for intended use will facilitate AD drug discovery and clinical trials through the enablement of precise selection of subjects at preclinical stages and evaluation of patients who are responsive vs nonresponsive to disease-modifying therapies.

To conclude, multicomponent biomarker (MCB) approaches[47, 48] have become increasingly important for their ability to characterize and dissect disease complexity (for example, diagnosis, staging, progression and stratification). Integrating omics-based biomarkers into MCB frameworks offers a powerful modality to increase precision and comprehensiveness in AD research and clinical applications.

## List of abbreviations

Aβ: Amyloid beta
AD: Alzheimer’s Disease
AUC: Area under the curve
CSF: Cerebrospinal fluid
DAM: Disease-associated microglia
DEG: Differentially expressed genes
DMT: Disease modifying therapy
GO: Gene ontology
GWAS: Genome-wide association study
MCI: Mild cognitive impairment
snRNA-seq: single nucleus RNA-seq
UMAP: Uniform manifold approximation and projection

## Declarations

### Ethics approval and consent to participate

The Religious Order Study (ROS) and Memory and Agins Project (MAP), collectively known as ROSMAP, adhere to a strict ethics statement centered on informed consent, participant autonomy and responsible data sharing. Both studies received approval from the Rush University Medical Center IRB and comply with the Declaration of Helsinki and local laws. All participants provided written informed consent and consented to the sharing of their de-identified data and biospecimens with qualified researchers through the RADC Sharing Hub and the AD Knowledge Portal, following IRB procedures. For the MCSA data, all study procedures and ethical aspects were reviewed and approved by Mayo Clinic’s Institutional Review Boards (IRBs) and Olmsted Medical Center. All participants were fully informed about the MCSA study’s scope and signed consent forms.

### Consent for publication

Not applicable.

### Availability of data and materials

The data used for this study are available from the Alzheimer’s Disease Knowledge Portal, for the single-nucleus RNAseq data, syn31512863; for the monocyte data, syn22024498; and for the MCI cohort, syn22024998.

All the code used for analysis is available at the GitHub repository, https://github.com/NCTrailRunner/microglia-biomarker

### Conflicts of Interest

Drs. Chiba-Falek and Lutz are coinventors of the related IP. Duke University filed a patent application for the technology developed in this study. Zhaohui Man, Yifei Zheng and Srilakshmi Venkatesan report no conflicts of interest.

### Funding

This work was funded in part by the National Institutes of Health/National Institute on Aging (NIH/NIA) RF1 AG077695 (OCF) and R01 AG057522 (OCF).

### Author Contributions

Conceptualization: OCF, MWL

Methodology: OCF, MWL, ZM

Investigation: OCF, MWL, ZM, SV, YZ

Visualization: OCF, MWL, ZM, YZ

Funding acquisition: OCF

Project administration: OCF

Supervision: OCF

Writing – original draft: OCF, MWL

Writing – review & editing: OCF, MWL

## Acknowledgments

The results published here are in whole or in part based on data obtained from the AD Knowledge Portal (https://adknowledgeportal.org). The data available in the AD Knowledge Portal would not be possible without the participation of research volunteers and the contribution of data by collaborating researchers.

Study data were provided by the Rush Alzheimer’s Disease Center, Rush University Medical Center, Chicago. Data collection was supported through funding from NIA grants P30AG10161 (ROS), R01AG15819 (ROSMAP; Genomics and RNA-seq), RC2AG036547 (H3K9Ac), R01AG36836 (RNA-seq), R01AG48015 (monocyte RNA-seq) RF1AG57473 (single nucleus RNA-seq), the Illinois Department of Public Health (ROSMAP), and the Translational Genomics Research Institute (genomic).

The snRNAseq data were generated from postmortem brain tissue provided by the Religious Orders Study and Rush Memory and Aging Project (ROSMAP) cohort at Rush Alzheimer’s Disease Center, Rush University Medical Center, Chicago. This work was funded by NIH grants U01AG061356 (De Jager/Bennett), RF1AG057473 (De Jager/Bennett), and U01AG046152 (De Jager/Bennett) as part of the AMP-AD consortium, as well as NIH grants R01AG066831 (Menon) and U01AG072572 (De Jager/St George-Hyslop).

