## Supplementary Methods for "A diagnostic plasma omics-biomarker for Alzheimer’s disease informed by microglial single-cell transcriptomics: A pilot study"

**Study design, data sources and quality control**

**For the ROSMAP post microglial samples, AD cases and control status were determined by consensus cognitive diagnosis. At the time of death, all available clinical data were reviewed by a neurologist with expertise in dementia, and a summary diagnostic opinion was rendered regarding the most likely clinical diagnosis at the time of death. Case conferences including one or more neurologists and a neuropsychologist were used for consensus on selected cases. AD cases were categorized as Alzheimer’s dementia (NINCDS probable or possible AD, cogdx = 4 or 5). Duration of Alzheimer’s disease ranged from 0 to 11 years (mean=1.8 years, SD=2.7).**

The quality control procedures excluded participants with mild cognitive impairment (cogdx = 2--3, 6) to focus on clearly delineated diagnostic categories (normal cognition vs Alzheimer’s disease). Missing cognitive diagnosis values in the source dataset were supplemented with postmortem neuropathological diagnosis (dcfdx_lv) when available, where dcfdx_lv represents standardized assessment of dementia status on the basis of brain tissue examination. After filtering, the microglial dataset comprised 270 samples (145 ADs with cogdx = 4--5, 125 normal samples with cogdx = 1), whereas the monocyte dataset included 236 samples (121 ADs, 115 cognitively normal samples). The monocyte samples were obtained

Replication cohort data were obtained from the Mayo Clinic study of Aging obtained from the Alzheimer’s Disease Knowledge Portal (syn22024998). The study included genetic and gene expression data from 422 participants. The generation was led by Dr. Nilufer Ertekin-Taner at the Mayo Clinic, Jacksonville, FL, as part of the multi-PI RF1AG051504 (MPIs Bu and Ertekin-Taner). This particular study focused on understanding pathomechanisms underlying cerebrovascular phenotypes that, in turn, may influence the risk of AD. Most of the individuals included in this study were cognitively unimpaired, a smaller proportion had MCI, and only 1 had AD.

**Transcriptomic data preprocessing**

The raw expression counts underwent standard preprocessing, including normalization to counts per million mapped reads, log₂ transformation with pseudocount addition, and filtering to retain genes expressed in at least 5% of the samples within each dataset. To enable cross-platform comparisons, we identified genes present in both datasets and applied variance stabilization. Binary diagnostic labels were assigned for model training, with cogdx = 1 classified as normal (0) and cogdx = 4--5 classified as AD (1).

**Biomarker gene panel construction**

We employed a systematic approach to identify optimal gene signatures by testing six complementary statistical feature selection strategies on the microglial training data. ANOVA and Student's t tests identified genes with the largest effect sizes between diagnostic groups. Coefficient of variation analysis revealed genes with high expression variability across samples. The limma-voom framework applies empirical Bayes moderation to improve statistical power by using information from all genes for variance estimation and is likely a better approach for small sample sizes, which is particularly valuable for smaller sample sizes. Random forest analysis provides importance scores reflecting each gene's contribution to classification accuracy through ensemble decision tree modeling. LASSO regression with elastic net penalties (α = 0.2) was used to perform automated feature selection while controlling for multicollinearity. Finally, the weighted composite score method combines normalized rankings from all other approaches via equal weighting. Some methods (Limma, ANOVA, t tests) prioritize the magnitude of genewise variance over effect size, whereas other methods adjust variance on the basis of the entire set of genes (LASSO). Random forest and lasso are also designed to capture nonlinear patterns in gene expression data.

For each selection strategy, we generated gene panels to systematically evaluate the relationship between feature set size and transfer learning performance. This approach enables the empirical determination of optimal panel configurations across different statistical frameworks.

**Deep Joint Distribution Optimal Transport architecture**

Cross-tissue label transfer uses a deep joint distribution optimal transport (DeepJDOT) neural network framework designed to align feature distributions between the source and target domains while preserving class-discriminative information. The architecture consisted of a shared encoder network with two fully connected layers (input → 2×hidden → hidden dimensions), incorporating batch normalization and dropout regularization (rates of 0.2--0.4) to prevent overfitting.

Classification employs an ordinal regression approach with a single threshold parameter, treating AD progression as an ordered categorical outcome rather than nominal classification. This design accounts for the natural ordering of cognitive states and has been shown to improve performance in medical classification tasks with inherent severity gradients.

**Training optimization and model selection**

The joint loss function combines standard cross-entropy classification loss on labeled microglial samples with the Sinkhorn-regularized optimal transport distance between microglial and monocyte feature embeddings. The Sinkhorn algorithm approximates the Wasserstein distance computation through iterative matrix scaling, enabling efficient gradient-based optimization. The regularization weights (λ_OT = 0.1--1.0) control the relative contribution of domain alignment versus classification accuracy.

Model training employs threefold stratified cross-validation to ensure balanced representation of diagnostic classes across folds. Hyperparameter optimization was used to evaluate the hidden layer dimensions (128, 256, 512), learning rates (10⁻³, 5×10⁻⁴, 10⁻⁴), and dropout rates (0.3, 0.4) via Cohen's kappa coefficient as the primary selection criterion to account for potential class imbalance. The F1 score and balanced accuracy served as secondary metrics.

To address class imbalance in the training data, we applied the synthetic minority oversampling technique (SMOTE) during model training and incorporated focal loss weighting to emphasize difficult-to-classify samples. Following hyperparameter selection, optimal models were retrained on the complete microglial dataset before application to monocyte samples.

**Label transfer evaluation and validation**

The trained models generated probabilistic predictions for each monocyte sample, which were subsequently converted to binary classifications via optimal probability thresholds determined via receiver operating characteristic analysis. The performance assessment included the accuracy, precision, recall, F1 score, and area under the ROC curve. We computed 95% confidence intervals for all the metrics via bootstrap resampling (n = 1000 iterations).

**UMAP visualization methods**

To generate integrated UMAP visualization of microglial and monocyte samples, we applied multistep batch correction and dimensionality reduction approaches. The expression data from both datasets were first normalized via SCTransform. The datasets were then subset to the selected gene panel (50-gene t test panel for the primary analysis, with the 30-gene combined panel available as an alternative). After identifying genes common to both datasets and removing dataset-specific genes with extreme expression differences (log_2_-fold change), we performed joint principal component analysis (PCA) on the combined expression matrix using the top 50 principal components.

To address batch effects between the brain microglia and peripheral blood monocyte datasets, we applied a two-step batch correction procedure: first, Harmony integration (theta=4, lambda=1.0, sigma=0.05, max iterations=50), followed by ComBat correction. Harmony integration was used for dimensionality reduction and ComBat was used to batch-correct the count matrices, allowing for valid differential expression and machine learning. The batch-corrected principal components were then used as inputs for UMAP dimensionality reduction via the UWOT R package with the following parameters: n_neighbors=70, min_dist=0.001, spread=3.0, metric="correlation", n_epochs=2000, and negative_sample_rate=20. These parameters were optimized to achieve clear separation of disease states while maintaining the integration of cell types across datasets.

**Regional Composition Analysis**

To quantify the spatial distribution of cell types and disease states across UMAP embedding, we performed a regional composition analysis. The UMAP coordinate space was partitioned into a 10×10 grid, and regions containing at least one sample were retained for analysis (n=35 occupied regions for the 50-gene panel; n=34 for the 30-gene panel). For each occupied region, we calculated the number and percentage of samples belonging to each of four categories: monocyte-normal, monocyte-AD, microglia-normal, and microglia-AD. The dominant category for each region was determined by identifying the category with the highest percentage of samples within that region. Regions were annotated with their region number, dominant cell type/disease category (Mon-N, Mon-AD, Mic-N, or Mic-AD), and corresponding percentage. This grid-based quantification enabled systematic assessment of cluster composition and identification of regions enriched for specific cell type and disease state combinations.

**Biological pathway enrichment analysis**

To understand the biological significance of the gene sets derived from differential expression analyses, we employed the Metascape program (Metascape.org)(1). Microglial gene sets were input as the target gene list, and all human genes were used for the background gene list. To group the top 20 enriched Metascape output GO terms into broader biological categories, the functions of each individual gene (based on a literature search) contributing to a particular category were qualitatively grouped into major functional categories. If a majority of the target genes associated with an enriched pathway GO term were associated with a particular functional category, this category was assigned to the GO term. A single GO term could be assigned to multiple functional categories. The minimum overlap of genes was 3, and a p value cutoff of 0.01 and minimum enrichment of 1.5 were used. For pathway databases, the GO biological processes, reactome gene sets, KEGG pathways and WikiPathways were used.

**Statistical analysis**

All computational analyses were performed via R version 4.3.0 and Python version 3.10 with standard scientific computing libraries. The cross-validation results are reported as the means ± standard deviations across folds. Statistical significance was assessed at α = 0.05 via appropriate tests for each comparison type. The weighted composite performance score incorporates several criteria with associated weights, such as Cohen's kappa (50%), the F1 score (35%), and accuracy (15%), with an emphasis on kappa to prioritize agreement beyond chance levels in this clinical prediction context.

1.	Zhou Y, Zhou B, Pache L, Chang M, Khodabakhshi AH, Tanaseichuk O, et al. Metascape provides a biologist-oriented resource for the analysis of systems-level datasets. Nat Commun. 2019;10(1):1523.
