## Extended Data Supplement for "A diagnostic plasma omics-biomarker for Alzheimer’s disease informed by microglial single-cell transcriptomics: A pilot study"

**Selection of gene panels of differentially expressed genes in microglia in AD**

The analysis utilized the microglial cell pseudobulk RNA-seq data extracted from this DLPFC snRNA-seq dataset and applied six different gene selection statistical methods: (1) ANOVA F statistics for multiclass comparisons or t tests for binary comparisons, (2) coefficient of variation (CV), (3) limma-based moderated F statistics, (4) random forest (RF) importance, (5) LASSO regression and (6) a weighted composite score based on the other algorithms. Each statistical method is optimal for specific characteristics of expression profiles and prioritizes different statistical properties (see Methods). Briefly, the t test captures genes with large effect sizes. The CV approach identifies the genes with the largest variance between the groups. The limma models utilize empirical Bayes statistics and are better for complex statistical designs with covariates. RF is a machine learning classifier that considers the level of contribution for each gene in terms of model accuracy. LASSO regression is a variant of linear and limma regression that provides a better signal-to-noise detection for a large number of genes. The combined method integrates multiple metrics (t test, CV, RF, limma, and lasso), creating a composite score.

Each method was applied to select the top panels of fewer than 100 genes—20--90 genes—for a total of 42 panels, with 7 per method. Similarly, the top panels with more than 100 genes (100--2000) were selected by each method to generate a total of 36 panels.

**Evaluation of the accuracy and precision of the different microglial gene panels in the classification of AD diagnosis**

For the 20-panel gene set, the LASSO and combined methods provided optimal F1 scores (48--52) and maintained larger scores over the range of panel sizes than the alternate methods did. The combined approach provided the highest F1 score (55) for the 30-gene panel. The t test showed a slightly better performance F1 score (57) than did all the other methods for panel sizes in the 40--70 gene range. The random forest, coefficient of variation and limma methods showed suboptimal performance according to the F1 score in contrast to the t test, combined and lasso methods for all panel sizes. For precision, the performance results largely followed F1 scores, with the t test showing the highest precision score (52) for panels of 30--70 genes. The precision of the lasso method is highest for the 20-gene panel, exceeding that of all other methods for this panel, but decreases steadily from 46 to 35 with increasing panel size. The random forest, coefficient of variation and limma methods achieved suboptimal performance in terms of precision compared with the other methods in terms of panel size. Interestingly, the combined score shows good performance (42--48) for small panel sizes of 20--30 genes but decreases to a level of 34 for panel sizes of 40 genes or more. The accuracy results largely correspond to the precision results. The best performance for panel sizes of 30--80 genes was for the t test (70% accuracy). The accuracy of LASSO was highest (65%) for the 20-gene panel and decreased to 40% with increasing gene panel size, up to 80 genes. The random forest, coefficient of variation and limma methods achieved suboptimal performance compared with the other methods in terms of accuracy. Recall is the proportion of actual positives (AD cases) that were correctly identified. For this measure, the combination method provides the best performance, with scores of 80--90 for panel sizes of 40--80 genes. The t test revealed recall levels of 60 consistently across panels of different sizes. LASSO improved the recall performance with increasing panel size, reaching a level of 90 for the 80 and 90 gene panels.

**Selected gene panels identified biochemical signaling pathways impaired in early-stage AD**

Biochemical pathway analysis was performed to identify coordinated biochemical signatures for the 300-gene panel. The enrichment analysis revealed the AD KEGG pathway (hsa05010) among the top 3 enriched pathways, as well as the actin-filament-based process and regulation of kinase activity (**Supplementary Figure 2**). The network of enriched biological pathways contained a set of highly connected pathways relevant to neuronal processes and neurodegeneration, including synapse organization, pathways related to neurodegeneration, chemokine signaling, axon extension and the regulation of kinase activity (**Supplementary Figure 3**). These pathways have been implicated in AD, and the results suggest that their involvement is in the early stages of the disease.

**Gene panel overlap with AD genome-wide association genes covers diverse cellular signaling pathways and functions**

We tested whether the genes selected for the gene panels were also found within AD-genome-wide association (GWAS) regions (i.e., associated SNPs +/- 500 kb) on the basis of the most recent GWAS[3]. In the 30-gene panel, we identified two GWAS genes: *ELDR,* an EGFR long noncoding RNA sequence, and *PTDGR,* a prostaglandin D2 receptor that responds to extracellular signals and activates intracellular signaling pathways. In the 50-gene panel, only one gene overlapped GWAS region, *SERPINE1*, a principal inhibitor of tissue plasminogen activator (tPA) and urokinase (uPA). For the 300-gene panel, we identified 13 genes, including *CELF2*, *PICALM*, *ELMO1*, *ATP8B4*, *SORL1*, *TRIO*, *UBE2K*, *LUC7L3*, *RBM47*, *RASA1*, *JAZF1*, *HERC1* and *CMIP*. This set of genes is involved in multiple signaling pathways, including RHO GTPases, RAS GTPases, and MAP kinases, which are critical for cytoskeletal remodeling and cellular signaling. These genes are also involved in endocytosis, RNA splicing and neuron projection development.

[1] Zeng L, Fujita M, Gao Z, White CC, Green GS, Habib N, et al. A Single-Nucleus Transcriptome-Wide Association Study Implicates Novel Genes in Depression Pathogenesis. Biol Psychiatry. 2023.

[2] Green GS, Fujita M, Yang HS, Taga M, McCabe C, Cain A, et al. Cellular dynamics across aged human brains uncover a multicellular cascade leading to Alzheimer's disease. bioRxiv. 2023.

[3] Bellenguez C, Kucukali F, Jansen IE, Kleineidam L, Moreno-Grau S, Amin N, et al. New insights into the genetic etiology of Alzheimer's disease and related dementias. Nat Genet. 2022;54:412–36.
