## Supplementary Figure 1 for "A diagnostic plasma omics-biomarker for Alzheimer’s disease informed by microglial single-cell transcriptomics: A pilot study"

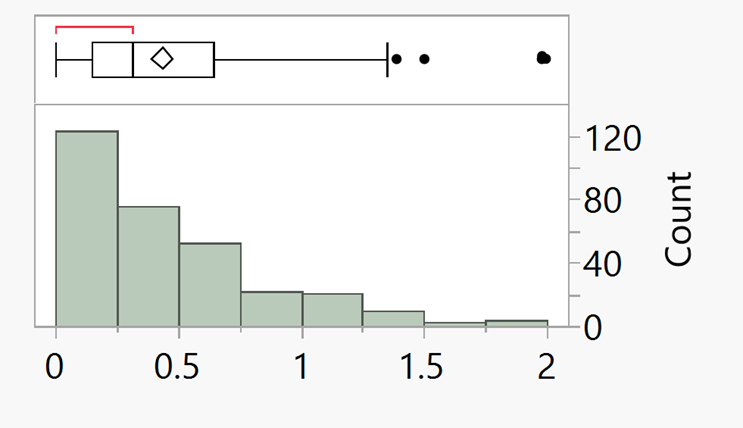

**Supplementary Figure 1. Distribution of -log10(P values) for the 300 gene panel selected by the limma algorithm.** The histogram shows the number of genes for each p value bucket. For the box plot, the vertical line shows the median, the confidence diamond contains the mean and upper and lower 95% of the mean, the ends of the box show the 25^th^ and 75^th^ quantiles, and the whiskers, extend from the box by 1.5*(interquartile range). The red bracket shows the densest 50% of all observations. Points outside the whiskers are considered outliers.
