## Supplementary Figure 2 for "A diagnostic plasma omics-biomarker for Alzheimer’s disease informed by microglial single-cell transcriptomics: A pilot study"

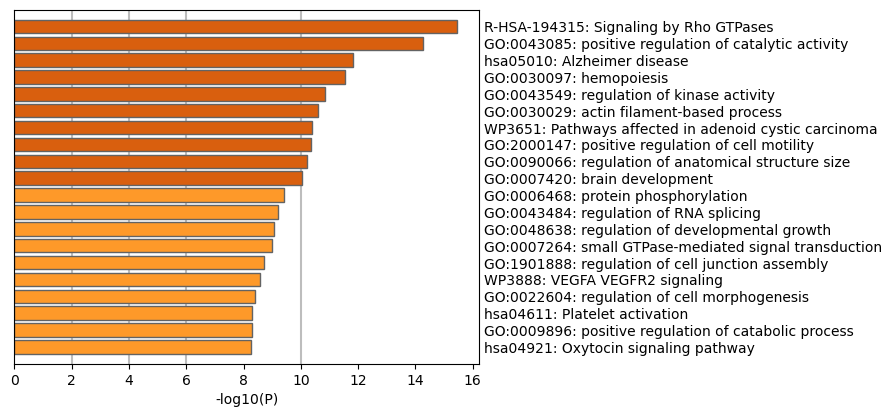

**Supplementary Figure 2. Metascape pathway analysis for 20 pathways identified by enrichment analysis for the 300 gene panel of microglial genes selected by limma.** The bar length corresponds to the -log of the p value for the enriched term (gene ontology).
