## Supplementary Figure 3 for "A diagnostic plasma omics-biomarker for Alzheimer’s disease informed by microglial single-cell transcriptomics: A pilot study"

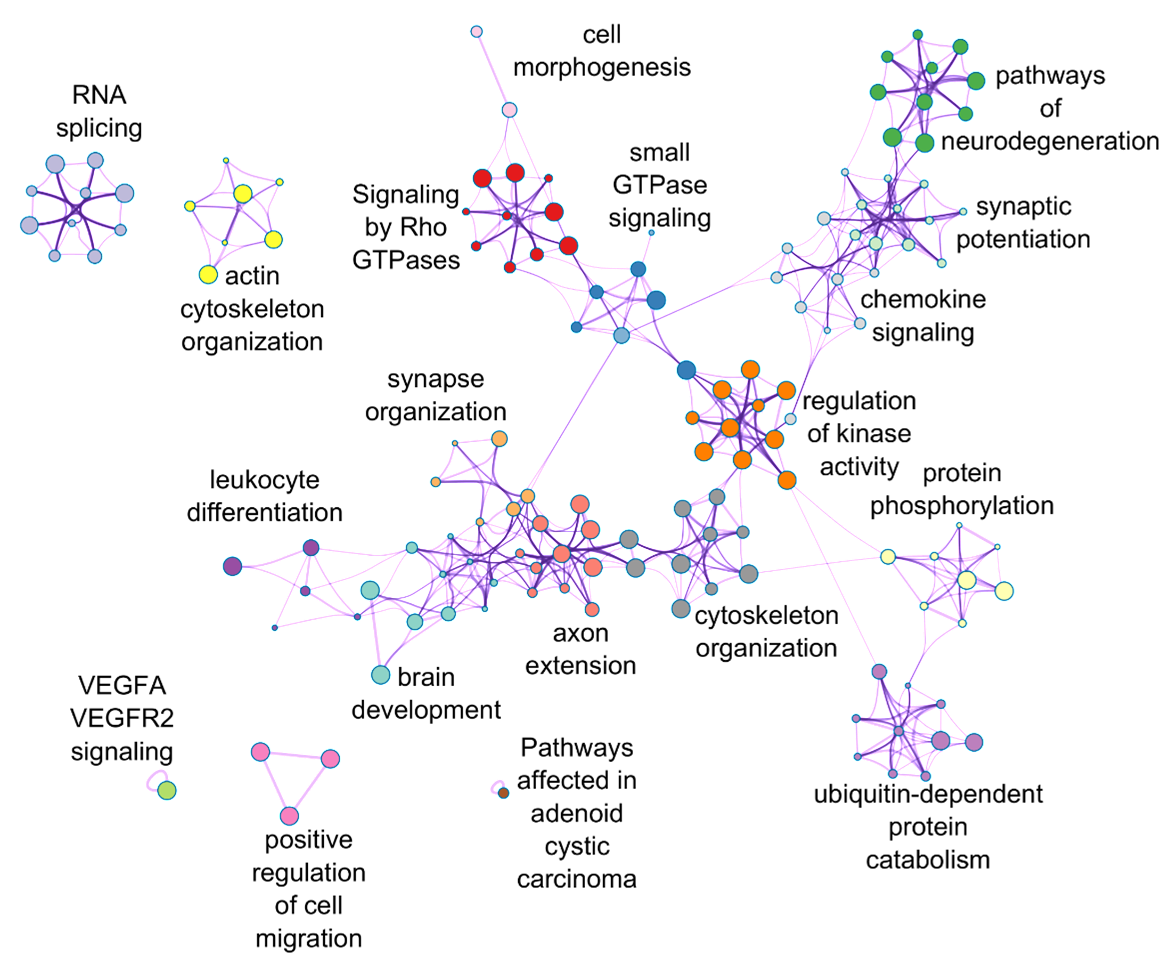

**Supplementary Figure 3.** **Enriched pathway network of the 300 DEGs in microglia.** Nodes represent groups of DEGs associated with individual biological pathways, linked by shared genes. Clusters of similar functions are labeled and color coded.
