## Supplemental Table S3 for "A diagnostic plasma omics-biomarker for Alzheimer’s disease informed by microglial single-cell transcriptomics: A pilot study"

**Supplemental Table S3. Performance measures for gene panels of different sizes for each statistical selection method.**

|  | Accuracy | | | | | |
| --- | --- | --- | --- | --- | --- | --- |
| **panel size** | **Combined** | **Lasso** | **Limma** | **Random Forest** | **t-test** | **Variance** |
| 20 | 60.17 | 65.68 | 30.51 | 35.59 | 61.86 | 44.49 |
| 30 | 66.74 | 58.90 |  | 40.04 | 68.64 | 44.07 |
| 40 | 37.71 | 54.45 | 33.05 | 51.06 | 67.58 | 32.63 |
| 50 | 44.49 | 51.69 |  | 45.34 | 69.07 | 58.26 |
| 70 | 46.19 | 44.70 | 29.66 | 45.97 | 68.64 | 55.30 |
| 80 | 45.55 | 38.77 |  | 58.69 | 55.72 | 45.55 |
| 90 | 31.78 | 41.53 | 42.37 | 52.75 | 63.98 | 54.24 |
| 100 | 40.04 | 41.10 | 31.57 | 44.49 | 67.16 | 51.06 |
| 300 | 43.64 | 36.02 | 59.75 | 39.62 | 37.50 | 40.25 |
| 500 | 47.88 | 60.17 | 44.92 | 47.67 | 37.29 | 36.65 |
| 800 | 32.63 | 58.05 |  | 32.20 | 40.68 | 67.58 |
| 1000 | 32.63 | 65.68 |  | 32.20 | 33.26 | 40.25 |
| 2000 | 51.48 | 38.77 |  | 32.84 | 38.56 | 31.78 |
|  | Recall | | | | | |
| **panel size** | **Combined** | **Lasso** | **Limma** | **Random Forest** | **t-test** | **Variance** |
| 20 | 0.67 | 0.49 | 0.42 | 0.34 | 0.36 | 0.41 |
| 30 | 0.64 | 0.73 |  | 0.27 | 0.57 | 0.68 |
| 40 | 0.94 | 0.70 | 0.71 | 0.19 | 0.61 | 0.94 |
| 50 | 0.83 | 0.69 |  | 0.41 | 0.65 | 0.19 |
| 70 | 0.83 | 0.76 | 0.48 | 0.35 | 0.65 | 0.25 |
| 80 | 0.80 | 0.90 |  | 0.27 | 0.55 | 0.68 |
| 90 | 0.76 | 0.89 | 0.90 | 0.27 | 0.64 | 0.61 |
| 100 | 0.51 | 0.92 | 0.58 | 0.48 | 0.08 | 0.51 |
| 300 | 0.87 | 0.95 | 0.75 | 0.29 | 0.91 | 0.42 |
| 500 | 0.86 | 0.69 | 0.23 | 0.21 | 0.97 | 0.96 |
| 800 | 0.99 | 0.71 |  | 0.67 | 0.83 | 0.05 |
| 1000 | 0.99 | 0.65 |  | 0.96 | 0.97 | 0.25 |
| 2000 | 0.79 | 0.87 |  | 1.00 | 0.93 | 0.97 |

|  | Precision | | | | | |
| --- | --- | --- | --- | --- | --- | --- |
| **panel size** | **Combined** | **Lasso** | **Limma** | **Random Forest** | **t-test** | **Variance** |
| 20 | 0.42 | 0.46 | 0.21 | 0.20 | 0.39 | 0.26 |
| 30 | 0.48 | 0.42 |  | 0.19 | 0.51 | 0.32 |
| 40 | 0.33 | 0.38 | 0.28 | 0.21 | 0.49 | 0.31 |
| 50 | 0.34 | 0.36 |  | 0.27 | 0.51 | 0.28 |
| 70 | 0.35 | 0.34 | 0.22 | 0.25 | 0.51 | 0.27 |
| 80 | 0.35 | 0.33 |  | 0.32 | 0.37 | 0.33 |
| 90 | 0.29 | 0.34 | 0.34 | 0.26 | 0.45 | 0.37 |
| 100 | 0.27 | 0.34 | 0.25 | 0.28 | 0.41 | 0.33 |
| 300 | 0.35 | 0.33 | 0.42 | 0.19 | 0.33 | 0.24 |
| 500 | 0.36 | 0.42 | 0.19 | 0.20 | 0.33 | 0.33 |
| 800 | 0.32 | 0.41 |  | 0.27 | 0.33 | 0.42 |
| 1000 | 0.32 | 0.47 |  | 0.31 | 0.32 | 0.18 |
| 2000 | 0.38 | 0.33 |  | 0.32 | 0.33 | 0.31 |
|  | F1 | | | | | |
| **panel size** | **Combined** | **Lasso** | **Limma** | **Random Forest** | **t-test** | **Variance** |
| 20 | 0.52 | 0.47 | 0.28 | 0.25 | 0.38 | 0.32 |
| 30 | 0.55 | 0.53 |  | 0.22 | 0.53 | 0.44 |
| 40 | 0.49 | 0.49 | 0.40 | 0.20 | 0.55 | 0.47 |
| 50 | 0.49 | 0.47 |  | 0.32 | 0.57 | 0.23 |
| 70 | 0.49 | 0.47 | 0.30 | 0.29 | 0.57 | 0.26 |
| 80 | 0.48 | 0.48 |  | 0.29 | 0.44 | 0.44 |
| 90 | 0.41 | 0.49 | 0.50 | 0.26 | 0.53 | 0.46 |
| 100 | 0.35 | 0.50 | 0.35 | 0.35 | 0.13 | 0.40 |
| 300 | 0.50 | 0.48 | 0.54 | 0.23 | 0.48 | 0.31 |
| 500 | 0.51 | 0.53 | 0.21 | 0.20 | 0.49 | 0.49 |
| 800 | 0.48 | 0.52 |  | 0.39 | 0.47 | 0.09 |
| 1000 | 0.48 | 0.54 |  | 0.47 | 0.48 | 0.21 |
| 2000 | 0.51 | 0.48 |  | 0.49 | 0.49 | 0.47 |
